# Adaptation of breast cancer cells to serum-free defined medium reveals changes in proliferation, adhesion, and lipid metabolism

**DOI:** 10.64898/2026.11.02.762606

**Authors:** Cassandra Aarrestad, Tala O. Khatib, James V. Alvarez

**Affiliations:** Translational Research Program, Public Health Sciences Division, Fred Hutchinson Cancer Center, Seattle, WA, United States of America

**Keywords:** serum-free media, defined media, breast cancer

## Abstract

Cancer cell metabolic dependencies are strongly influenced by nutrient availability, yet conventional culture conditions contain non-physiological nutrient concentrations and poorly defined serum components that may obscure metabolic vulnerabilities. Here, we investigated the adaptation of three breast cancer cell lines to physiologically relevant, serum-free defined medium (DM). Using a stepwise reduction in fetal bovine serum (FBS), we evaluated proliferation, adhesion, morphology, and transcriptional changes in HER2-positive SK-BR-3, triple-negative MDA-MB-231, and ER-positive/HER2-positive BT-474 cells. The three cell lines differed substantially in their ability to adapt to serum-free conditions. MDA-MB-231 cells adapted readily, maintaining proliferation and adhesion without exogenous substrate coatings. SK-BR-3 cells also achieved sustained proliferation in serum-free DM, but required a serum-derived coating to maintain adhesion. In contrast, BT-474 cells failed to sustain adhesion and proliferation following complete serum withdrawal, although they remained proliferative in medium containing 1% FBS. Adapted cells retained characteristic HER2 or vimentin expression despite changes in morphology and proliferation. RNA sequencing revealed extensive transcriptional remodeling, including reduced expression of cell-cycle-associated programs in SK-BR-3 cells and enrichment of cholesterol homeostasis gene sets in SK-BR-3 and MDA-MB-231 cells. Expression of cholesterol regulatory genes, including *SREBF2*, *DHCR24,* and *FDFT1*, increased across all three lines. Together, these findings highlight cell-line-specific adhesion requirements and shared metabolic responses to serum withdrawal, establishing a framework for developing defined culture systems to investigate cancer metabolic dependencies through genetic screening.

## Introduction

Metabolic reprogramming is a hallmark of cancer that enables tumor cells to sustain proliferation and survival under changing environmental conditions [1–4]. The metabolic requirements of cancer cells are shaped not only by intrinsic oncogenic programs but also by the availability of nutrients, oxygen, and other extracellular factors [3,4]. Consequently, the metabolic behavior of cancer cells can be highly sensitive to their environment. Conventional cell culture media, however, was developed primarily to support robust cellular proliferation rather than to reproduce the nutrient composition of human tissues or plasma. Many commonly used formulations contain concentrations of nutrients and ions that differ substantially from those encountered in vivo. For example, RPMI 1640 contains approximately 0.4 mM calcium, compared with approximately 2–2.5 mM total calcium in human plasma [5]. Such differences can alter cellular metabolism, signaling, and proliferation, potentially limiting the physiological relevance of findings obtained using conventional culture conditions.

To address these limitations, several cell culture media formulations have been developed to more closely approximate the metabolic composition of human plasma. Human plasma-like medium (HPLM), for example, contains approximately 60 components at concentrations based on those measured in human plasma [6]. Compared with conventional media, HPLM leads to changes in intracellular metabolite pools, redox state, and glucose utilization in cancer cells [6]. Additional studies have demonstrated that physiological media influence signaling pathways, gene essentiality, and broader transcriptional and proteomic programs [7,8]. These effects extend beyond cancer metabolism, with physiological media influencing cellular maturation, immune activation, and responses to environmental stimuli in diverse experimental models [5,9,10]. Similar findings have been reported using Plasmax, another medium designed to approximate physiological nutrient concentrations, which alters proliferation, migration, and glucose consumption in breast cancer cells [11]. Collectively, these studies demonstrate that nutrient composition is an important determinant of cellular phenotypes and highlight the potential limitations of conventional culture media.

Despite these advances, an important source of variability in cell culture remains: supplementation with fetal bovine serum (FBS). Serum is widely used to support cell growth and survival but introduces a complex mixture of proteins, growth factors, hormones, lipids, metabolites, and other components whose concentrations are often poorly characterized [12,13]. Furthermore, FBS composition varies between production lots, potentially contributing to experimental variability and limiting reproducibility [14–16]. Importantly, the use of physiological basal media does not necessarily eliminate these concerns, as many studies continue to supplement these formulations with serum, including dialyzed or reduced-serum preparations [6–8,11]. Thus, although physiological media offer greater control over nutrient availability, serum supplementation introduces additional biological factors that remain incompletely defined.

Eliminating serum presents particular challenges for adherent cells, which depend on extracellular signals for attachment, survival, and proliferation. In addition to soluble growth factors, serum contains extracellular matrix proteins, including fibronectin and vitronectin, that can adsorb to culture surfaces and promote cell adhesion [17,18]. Serum withdrawal can therefore disrupt cellular attachment and morphology, complicating efforts to establish serum-free culture conditions [12,13]. Although chemically defined and serum-free media hav7e been developed for numerous cell types [19–24], adapting adherent cancer cells to these conditions often requires additional support for attachment and growth [13,18]. Previous studies have demonstrated that MDA-MB-231 breast cancer cells can be maintained in serum-free formulations when cultured on extracellular matrix-coated surfaces [25,26]. However, these approaches have not fully established whether breast cancer cells with distinct molecular characteristics can be maintained under serum-free conditions with physiologically relevant nutrient concentrations. Furthermore, the cellular and transcriptional changes accompanying adaptation to these conditions remain incompletely understood.

Here, we investigated the feasibility of adapting three breast cancer cell lines with distinct molecular characteristics – HER2-positive SK-BR-3 cells, triple-negative MDA-MB-231 cells, and ER-positive/HER2-positive BT-474 cells – to a defined medium (DM) without serum supplementation. Using a stepwise reduction in FBS concentration and substrate coatings to support cellular attachment, we evaluated cell proliferation, morphology, and transcriptional responses during adaptation. SK-BR-3 and MDA-MB-231 cells were successfully maintained in serum-free DM, whereas BT-474 cells failed to sustain attachment following complete serum withdrawal and were instead characterized under reduced-serum conditions. Transcriptomic analysis revealed downregulation of cell-cycle-associated programs and increased expression of cholesterol-homeostasis genes across all three adapted populations. Together, these findings establish a framework for culturing breast cancer cells under more defined nutritional conditions and reveal shared and cell-line-specific responses to reduced serum availability.

## Results

### Stepwise serum withdrawal enables adaptation of SK-BR-3 cells to serum-free defined medium

To culture breast cancer cells under conditions that more closely approximate the nutrient composition of the tumor microenvironment, we used a defined medium (DM) using human plasma-like medium (HPLM), adapted from previously described defined formulations [27–29]. HPLM provides metabolites at concentrations that more closely resemble those found in human plasma than conventional culture media. The DM was supplemented with insulin-transferrin-selenium-ethanolamine (ITSE) to provide insulin signaling and essential micronutrients; hydrocortisone to support epithelial cell growth and survival; and human epidermal growth factor (EGF) to provide mitogenic signaling. Bovine serum albumin (BSA) was included as a carrier protein, and oleic and palmitic acid were added to provide physiologically relevant sources of unsaturated and saturated fatty acids, respectively.

Direct transfer of breast cancer cells from medium containing 10% FBS into serum-free DM resulted in minimal proliferation (data not shown). Because gradual serum withdrawal has been used to adapt mammalian cells to chemically defined or serum-free culture conditions [21,22], we developed a stepwise adaptation protocol in which complete medium (CM; containing 10% FBS) was progressively replaced with DM (Figure 1A). Throughout this study, media compositions are expressed as CM:DM ratios. For example, a 50:50 CM:DM ratio corresponds to a final FBS concentration of 5%.

**Figure 1.**
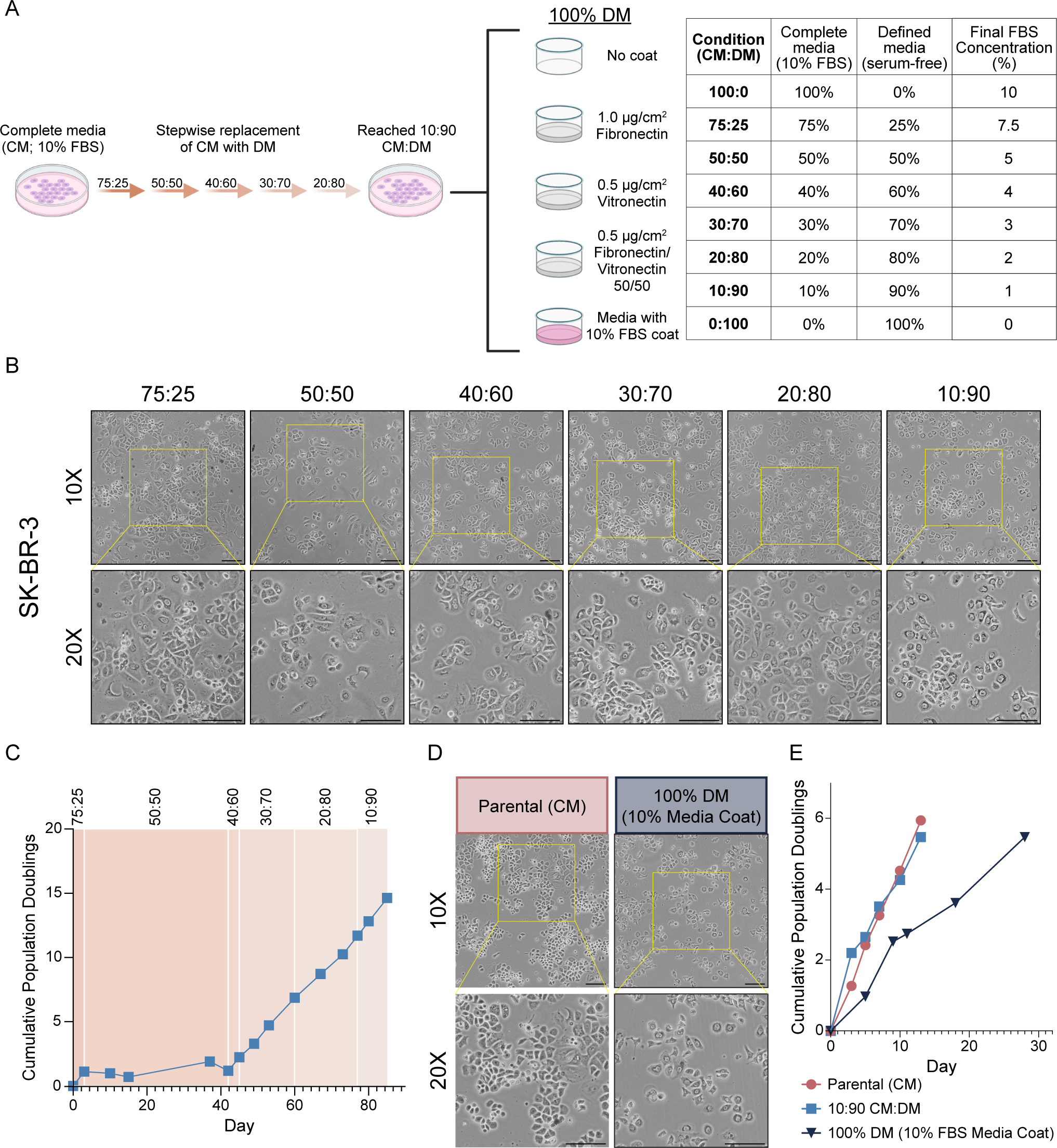
Stepwise serum withdrawal and a serum-derived plate coating enable SK-BR-3 adaptation to serum-free defined medium. **(A)** Schematic of the stepwise adaptation protocol used to transition SK-BR-3 cells from complete medium containing 10% FBS (CM) to serum-free defined medium (DM), and of the plate-coating conditions tested at the 100% DM endpoint. Media compositions are expressed as CM:DM ratios, where CM is complete medium containing 10% FBS and DM is serum-free defined medium. **(B)** Representative phase-contrast images of SK-BR-3 cells at each stage of adaptation. **(C)** Cumulative population doublings during stepwise adaptation. **(D)** Representative phase-contrast images of parental SK-BR-3 cells in CM and adapted cells maintained in 100% DM on plates pre-coated with medium containing 10% FBS. **(E)** Cumulative population doublings of parental cells, cells at the 10:90 adaptation stage, and cells maintained in 100% DM on serum-coated plates. Scale bars, 150 µm. Panel (A) was created in BioRender and assembled with additional labels and layout in Adobe Illustrator. Created in BioRender. Aarrestad, C. (2026) https://BioRender.com/okp0u7s.

We first characterized the response of SK-BR-3 cells during this adaptation process. Cell morphology changed modestly as the proportion of DM increased, suggesting that the cells underwent phenotypic changes during adaptation (Figure 1B). These changes were accompanied by a transient decrease in proliferation. At a CM:DM ratio of 50:50 (5% FBS), cells proliferated at only 0.0017 population doublings/day. However, proliferation increased to 0.313 population doublings/day after transition to a CM:DM ratio of 40:60 (4% FBS; Figure 1C). Thus, the marked reduction in proliferation at the 50:50 stage was transient and was followed by recovery as cells continued through the adaptation protocol.

In addition to providing nutrient support, serum also provides proteins that enable cell attachment to tissue culture plates. Consistent with this, as the cells approached complete serum withdrawal, serum was required for sustained adhesion under standard culture conditions. We therefore compared plates coated with fibronectin (FN); vitronectin (VN); a combination of FN and VN; 10% FBS-containing medium; or no coating. SK-BR-3 cells initially attached under all conditions; however, only cells plated on surfaces pre-coated with 10% FBS-containing medium remained adherent after passaging. Cells cultured on FN-, VN-, or FN/VN-coated plates, as well as uncoated plates, progressively detached and were not maintained with continued culture (Supplementary Figure 1). We therefore used the serum-derived coating for subsequent culture of cells in 100% DM.

Fully adapted SK-BR-3 cells maintained in 100% DM on serum-coated plates remained epithelial in appearance but displayed altered colony organization relative to parental cells, forming less densely packed and more dispersed clusters (Figure 1D). Cells maintained at a CM:DM ratio of 10:90 (1% FBS) proliferated at a rate similar to parental SK-BR-3 cells cultured in CM (0.42 versus 0.46 population doublings/day, respectively). In contrast, proliferation decreased to 0.25 population doublings/day following transition to 100% DM (Figure 1E). Despite this reduction, cells maintained consistent proliferation across five passages in serum-free DM. Together, these results establish a stepwise method for adapting SK-BR-3 cells to serum-free DM and show that the adapted cells can be maintained long-term, although with reduced proliferation and altered colony organization relative to cells grown under conventional serum-containing conditions.

### MDA-MB-231 cells readily adapt to defined medium, whereas BT-474 cells fail to maintain adhesion under serum-free conditions

Having successfully adapted HER2+ SK-BR-3 cells to DM, we next investigated whether the same approach could be applied to breast cancer cell lines representing other molecular subtypes. We selected triple-negative MDA-MB-231 cells and luminal B BT-474 cells to evaluate their ability to proliferate and maintain adhesion following serum withdrawal.

Of the three cell lines examined, MDA-MB-231 cells exhibited the most robust adaptation to DM. Unlike SK-BR-3 cells, MDA-MB-231 cells did not undergo a discernible proliferative bottleneck during adaptation, maintaining continuous growth as CM was progressively replaced with DM (Figure 2A). Following transition to 100% DM, MDA-MB-231 cells remained adherent and proliferative across all extracellular matrix (ECM) coating conditions tested, including uncoated plates, throughout five serial passages (Supplementary Figure 1). Cells cultured on VN-coated plates achieved the greatest cumulative population doublings (cf. Figure 2C and Supplementary Figure 1D), and this condition was therefore selected for subsequent analyses.

**Figure 2.**
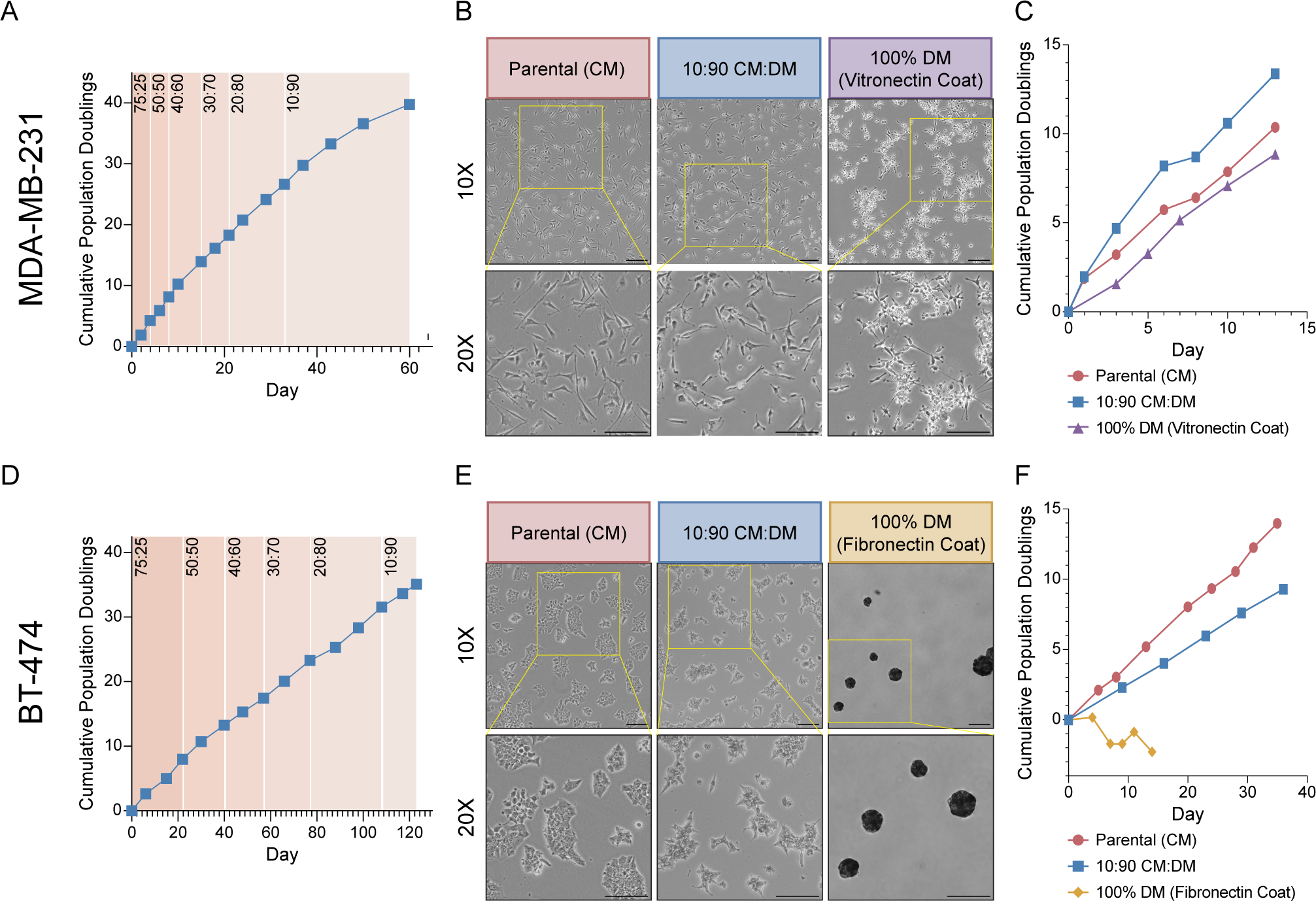
MDA-MB-231 and BT-474 cells differ in their ability to adapt to serum-free defined medium. **(A)** Cumulative population doublings of MDA-MB-231 cells during stepwise adaptation to DM. **(B)** Representative phase-contrast images of parental MDA-MB-231 cells maintained in CM, cells at the 10:90 CM:DM adaptation stage, and cells maintained in 100% DM on VN-coated plates. **(C)** Cumulative population doublings of parental, 10:90, and adapted 100% DM MDA-MB-231 cultures. **(D)** Cumulative population doublings of BT-474 cells during stepwise adaptation. **(E)** Representative phase-contrast images of parental BT-474 cells maintained in CM, cells at the 10:90 CM:DM adaptation stage, and cells transferred to 100% DM on FN-coated plates. **(F)** Cumulative population doublings of parental, 10:90, and 100% DM BT-474 cultures. Scale bars, 150 µm.

Parental MDA-MB-231 cells displayed the characteristic dispersed, mesenchymal morphology of this cell line, which was largely maintained at a CM:DM ratio of 10:90 (1% FBS). In contrast, cells cultured in 100% DM on VN-coated plates formed more compact clusters while retaining elongated cellular protrusions characteristic of a mesenchymal phenotype (Figure 2B). MDA-MB-231 cells maintained at a CM:DM ratio of 10:90 proliferated more rapidly than parental cells cultured in CM, exhibiting growth rates of 1.03 and 0.80 population doublings/day, respectively (Figure 2C). Although proliferation decreased to 0.68 population doublings/day following transition to 100% DM, cells maintained sustained growth in the absence of serum. Together, these findings demonstrate that MDA-MB-231 cells readily adapt to serum-free DM while maintaining adhesion and proliferation.

In contrast, BT-474 cells exhibited greater sensitivity to serum withdrawal. BT-474 cells are an estrogen receptor-positive (ER+), luminal B breast cancer cell line characterized by relatively slow proliferation and the formation of compact epithelial clusters. To compensate for the removal of serum-derived estradiol [30], exogenous estradiol was added beginning at a CM:DM ratio of 40:60 (4% FBS). Although BT-474 cells did not exhibit the pronounced proliferative bottleneck observed in SK-BR-3 cells, their growth declined during certain stages of adaptation. This was most evident following transition to a CM:DM ratio of 20:80 (2% FBS), during which cells proliferated at 0.10 population doublings/day, compared with an overall proliferation rate of 0.29 population doublings/day during adaptation (Figure 2D). Proliferation subsequently recovered following transition to the 10:90 condition, indicating that BT-474 cells could tolerate substantial serum reduction.

However, BT-474 cells failed to maintain sustained adhesion and proliferation following complete serum withdrawal. Although cells initially adhered to all coating conditions tested in 100% DM, they detached following the first passage, with progressive loss of viable cells during subsequent passages (Supplementary Figure 1). FN-coated plates supported the greatest cumulative population doublings (Figure 2F), although none of the coating conditions permitted sustained growth.

Morphological analysis revealed that parental BT-474 cells formed densely packed epithelial clusters, whereas cells maintained at a CM:DM ratio of 10:90 formed smaller, more dispersed clusters while retaining an epithelial appearance. In contrast, BT-474 cells cultured in 100% DM on FN-coated plates detached from the substrate and formed large aggregates in suspension (Figure 2E). Consistent with these observations, proliferation declined with progressive serum withdrawal. Parental BT-474 cells cultured in CM proliferated at 0.40 population doublings/day, compared with 0.26 population doublings/day at a CM:DM ratio of 10:90 (Figure 2F). In 100% DM, the population doubling rate declined further to −0.16/day, consistent with a net loss of cells following detachment.

Collectively, these findings reveal substantial differences in the ability of breast cancer cell lines to adapt to serum-free DM. Whereas SK-BR-3 cells required an extended adaptation period and MDA-MB-231 cells adapted with minimal difficulty, BT-474 cells failed to maintain long-term adhesion and proliferation in serum-free DM under the coating conditions tested. Nevertheless, BT-474 cells could be maintained under substantially reduced-serum conditions, demonstrating that partial serum withdrawal is tolerated even when complete adaptation is unsuccessful.

### Breast cancer cells retain characteristic phenotypic markers following adaptation to reduced-serum or serum-free defined medium

Having successfully adapted SK-BR-3 and MDA-MB-231 cells to serum-free defined medium (DM), and BT-474 cells to medium containing 1% fetal bovine serum (FBS), we next investigated whether adaptation was associated with changes in cellular phenotype. To assess whether adapted cells retained characteristic molecular features, we examined HER2 expression in SK-BR-3 and BT-474 cells and vimentin expression in MDA-MB-231 cells. We also evaluated the proliferation marker Ki-67 to determine how adaptation influenced the proportion of cycling cells (Figure 3).

**Figure 3.**
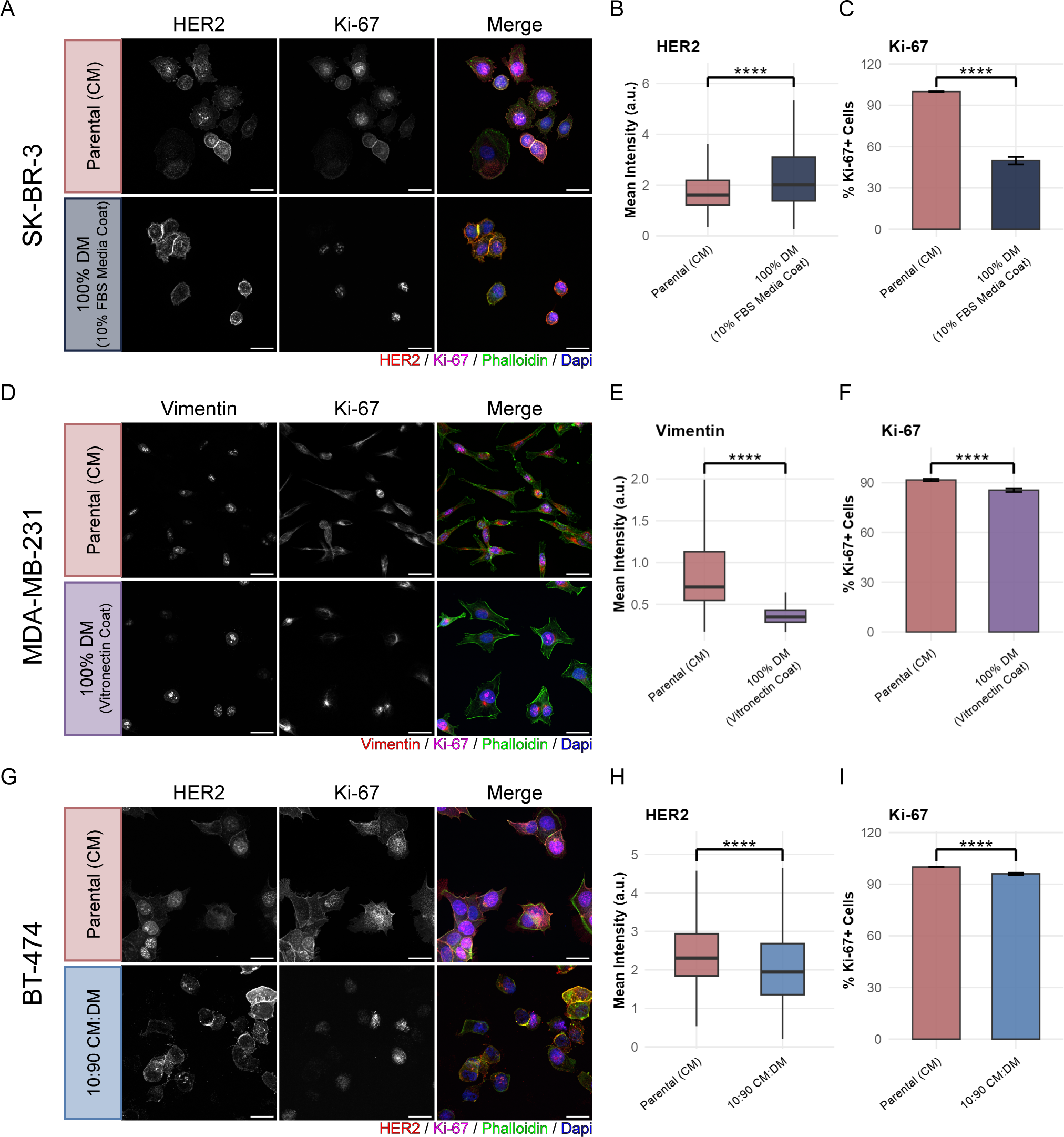
Adapted breast cancer cells retain subtype-associated markers while remaining proliferative in defined medium. **(A)** Representative immunofluorescence images of HER2 and Ki-67 in parental SK-BR-3 cells and adapted cells maintained in 100% DM on serum-coated plates. **(B)** Quantification of HER2 mean fluorescence intensity. **(C)** Percentage of Ki-67-positive cells. **(D–F)** Representative immunofluorescence images of vimentin and Ki-67 **(D)**, quantification of vimentin mean fluorescence intensity **(E)**, and percentage of Ki-67-positive cells **(F)** in parental and adapted MDA-MB-231 cells maintained in 100% DM. **(G–I)** Representative immunofluorescence images of HER2 and Ki-67 **(G)**, quantification of HER2 mean fluorescence intensity **(H)**, and percentage of Ki-67-positive cells **(I)** in parental BT-474 cells and cells maintained at the 10:90 CM:DM adaptation stage. Phalloidin and DAPI label F-actin and nuclei, respectively. Representative images were acquired using a 63× objective; quantitative measurements were obtained from 20× montage images. Cell numbers quantified for SK-BR-3: n = 1,394 for parental and n = 333 for adapted; for MDA-MB-231: n = 1,786 for parental and n = 1,054 for adapted; and for BT-474: n = 2,729 and n = 1,193. Box plots show the median and interquartile range, with whiskers extending to 1.5× the interquartile range. Bar graphs show the mean ± SEM. ****P < 0.0001, two-sided Wilcoxon rank-sum test with Benjamini-Hochberg correction. Antibody and reagent information is provided in Table S2.

Because SK-BR-3 cells underwent a pronounced proliferative bottleneck during adaptation, we first investigated whether the resulting population retained its characteristic HER2-positive phenotype. Immunofluorescence imaging revealed robust HER2 expression in both parental and adapted cells (Figure 3A). Quantification showed a modest increase in mean HER2 fluorescence intensity following adaptation to 100% DM, accompanied by greater variability in expression among individual cells (Figure 3B). In contrast, the proportion of Ki-67-positive cells was significantly reduced in the adapted population (p < 0.0001), consistent with the slower proliferation observed in serum-free culture (Figure 3C). Thus, SK-BR-3 cells retained HER2 expression following adaptation despite a marked reduction in proliferation.

In MDA-MB-231 cells, adaptation to serum-free DM was accompanied by subtle morphological changes. Although adapted cells retained an elongated, mesenchymal appearance, their cellular protrusions were broader and less extended than those of parental cells (Figure 3D). These changes were accompanied by a significant reduction in vimentin fluorescence intensity (p < 0.0001; Figure 3E). The proportion of Ki-67-positive cells was also reduced following adaptation, although a substantial population of cells remained Ki-67-positive (Figure 3F). Together, these findings indicate that MDA-MB-231 cells retain mesenchymal features following adaptation to serum-free DM, despite changes in morphology, vimentin expression, and proliferative activity.

Because BT-474 cells failed to maintain adhesion and proliferation in serum-free DM, we characterized cells maintained under the 10:90 medium condition containing 1% FBS. Immunofluorescence imaging revealed that these cells retained HER2 expression, although mean fluorescence intensity was modestly reduced compared with parental cells (Figure 3G,H). The proportion of Ki-67-positive cells was also lower following adaptation, consistent with the reduced proliferation observed under these conditions (Figure 3I). Nevertheless, the adapted population remained HER2-positive and contained proliferating cells despite the substantial reduction in serum concentration.

Collectively, these results demonstrate that adaptation to serum-free or reduced-serum DM preserves key phenotypic characteristics of the breast cancer cell lines examined, although adaptation is accompanied by changes in cell morphology, marker expression, and the proportion of proliferating cells.

### Adaptation to defined medium induces distinct transcriptional programs associated with cell-cycle regulation and metabolic remodeling

Having established that adapted breast cancer cells retained characteristic phenotypic markers despite changes in morphology and proliferation, we next investigated the transcriptional consequences of adaptation using bulk RNA sequencing (RNA-seq). We compared parental cells maintained in CM with cells adapted to 100% DM (SK-BR-3 and MDA-MB-231) or a CM:DM ratio of 10:90 (BT-474), performed differential gene expression analysis using DESeq2, and identified differentially expressed genes using an adjusted p-value below 0.05 and an absolute log2 fold change greater than 1. To identify biological pathways associated with adaptation, we performed gene set enrichment analysis (GSEA) using the MSigDB Hallmark gene sets (Figure 4).

**Figure 4.**
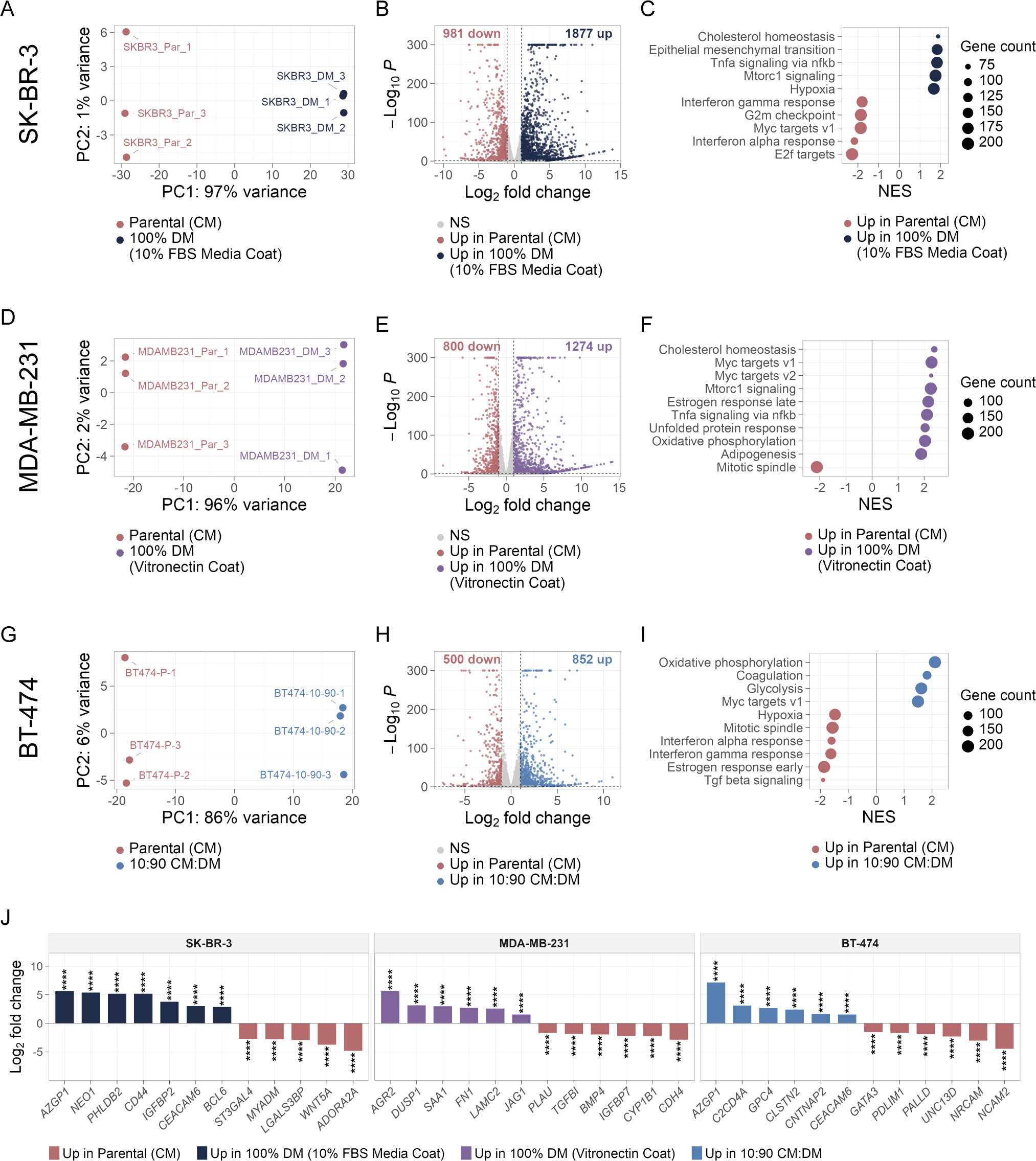
Adaptation to defined medium produces shared and cell-line-specific transcriptional changes. **(A, D, G)** Principal component analysis of parental and adapted SK-BR-3, MDA-MB-231, and BT-474 cells, respectively. **(B, E, H)** Volcano plots showing differential gene expression in adapted relative to parental cells. Differentially expressed genes were defined by an adjusted P value < 0.05 and an absolute log, fold change > 1; genes meeting these criteria are colored according to the direction of change. **(C, F, I)** Gene set enrichment analysis using Hallmark gene sets in SK-BR-3, MDA-MB-231, and BT-474 cells, respectively. Positive normalized enrichment scores (NES) indicate enrichment in adapted cells, whereas negative NES indicate enrichment in parental cells; dot size represents the number of genes contributing to each gene set. SK-BR-3 and MDA-MB-231 cells were analyzed following adaptation to 100% DM, whereas BT-474 cells were analyzed at the 10:90 CM:DM adaptation stage (1% FBS).

Among the three cell lines, SK-BR-3 cells exhibited the greatest transcriptional separation between parental and adapted conditions. Principal component analysis (PCA) distinguished the two populations along the first principal component (PC1), which accounted for 98% of the variance (Figure 4A). Differential expression analysis identified 1,877 upregulated and 981 downregulated genes in cells adapted to 100% DM relative to parental cells maintained in CM (Figure 4B). GSEA revealed reduced expression of genes associated with cell-cycle progression, including E2F targets (NES = −2.31), G2M checkpoint (NES = −1.90), and MYC targets V1 (NES = −1.90), consistent with the reduced proliferation observed following adaptation. Interferon-α and interferon-γ response gene sets were also enriched in parental cells. In contrast, adapted cells exhibited enrichment of cholesterol homeostasis (NES = 1.82), TNFα signaling via NF-κB (NES = 1.79), epithelial–mesenchymal transition (NES = 1.79), mTORC1 signaling (NES = 1.69), and hypoxia (NES = 1.65) gene sets (Figure 4C). These findings suggest that adaptation in SK-BR-3 cells is associated with reduced cell-cycle activity and broad changes in metabolic and stress-associated transcriptional programs.

MDA-MB-231 cells also exhibited substantial transcriptional changes following adaptation to serum-free DM, with parental and adapted samples separating along PC1, which explained 95% of the variance (Figure 4D). Differential expression analysis identified 1,274 upregulated and 800 downregulated genes in adapted cells (Figure 4E). In contrast to SK-BR-3 cells, the transcriptional response of MDA-MB-231 cells was characterized primarily by enrichment of metabolic and growth-associated pathways. Cholesterol homeostasis was the most strongly enriched Hallmark gene set (NES = 2.28), accompanied by enrichment of MYC targets, mTORC1 signaling, oxidative phosphorylation, and adipogenesis. In contrast, the mitotic spindle gene set was enriched in parental cells (NES = −2.06; Figure 4F). Thus, adaptation of MDA-MB-231 cells was associated with broad metabolic remodeling, particularly in pathways related to cholesterol homeostasis, alongside changes in cell-cycle-associated gene expression.

Although BT-474 cells could not sustain growth in 100% DM, adaptation to a CM:DM ratio of 10:90 (1% FBS) was nevertheless accompanied by substantial transcriptional changes. PCA distinguished parental cells maintained in CM from cells adapted to 10:90 CM:DM along PC1, which accounted for 86% of the variance (Figure 4G). Differential expression analysis identified 852 upregulated and 500 downregulated genes in adapted cells (Figure 4H). GSEA revealed enrichment of oxidative phosphorylation (NES = 2.07), coagulation (NES = 1.80), glycolysis (NES = 1.60), and MYC targets V1 (NES = 1.51) in adapted cells. Conversely, estrogen response, TGF-β signaling, mitotic spindle, and interferon response gene sets were enriched in parental cells (Figure 4I). These findings indicate that even partial serum withdrawal is associated with substantial transcriptional remodeling in BT-474 cells, including changes in metabolic and proliferation-associated pathways.

To determine whether the different adhesion behaviors observed during adaptation were reflected at the transcript level, we analyzed GO Biological Process gene sets with “adhesion” in their name using the same DESeq2 pipeline applied to the Hallmark sets (Supplementary Table S1). Adhesion-associated genes were differentially expressed in all three lines, but the specific genes that changed differed between lines (Figure 4J). AZGP1 and CEACAM6 were upregulated in all three adapted populations (log2 fold change relative to parental cells = 5.6, 10.6, and 7.1 for AZGP1 and 3.0, 6.7, and 1.6 for CEACAM6, in SK-BR-3, MDA-MB-231, and BT-474 cells, respectively). Each line also showed distinct changes: adapted SK-BR-3 cells upregulated CD44 (5.2) and downregulated ADORA2A (−4.8); adapted MDA-MB-231 cells upregulated AGR2 (5.6), FN1 (2.7), and LAMC2 (2.6) and downregulated CDH4 (−2.8); and adapted BT-474 cells downregulated NCAM2 (−4.5) and NRCAM (−3.0). WNT5A changed in opposite directions, downregulated in SK-BR-3 cells (−3.8) but upregulated in MDA-MB-231 cells (2.3).

Collectively, these analyses demonstrate that adaptation to defined medium, either fully or through partial replacement of CM, induces extensive, cell-line-dependent transcriptional changes. Although the specific responses differ among cell lines, alterations in metabolic and cell-cycle-associated programs emerge as recurring features of adaptation, and the adhesion-associated genes that changed differed in each line.

## Discussion

The results of this study demonstrate that breast cancer cells can be adapted to growth in serum-free, physiologically inspired defined medium (DM), although the ability to establish and maintain proliferation under these conditions varies substantially among cell lines. MDA-MB-231 cells adapted relatively readily, maintaining proliferation throughout the adaptation process and remaining adherent in 100% DM without exogenous extracellular matrix (ECM) coatings. In contrast, SK-BR-3 cells underwent a prolonged adaptation period characterized by transient reductions in proliferation and required a serum-derived substrate coating to maintain adhesion. BT-474 cells tolerated substantial serum reduction but failed to maintain sustained growth in 100% DM, even in the presence of ECM coatings. These differences are consistent with previous reports that the ability to adapt to serum-free culture depends on cell type and medium composition [21]. Interestingly, the relative ease with which MDA-MB-231 cells adapted may be related to their mesenchymal phenotype, as EMT-like breast cancer cells have been reported to exhibit enhanced survival under serum-free conditions [31]. However, additional cell lines will be required to determine whether mesenchymal characteristics or molecular subtype are predictive of adaptation capacity.

An important finding of this study is that serum withdrawal introduces challenges beyond nutrient availability, particularly with respect to cell adhesion. In addition to supplying nutrients and growth factors, serum contains proteins that facilitate attachment to culture surfaces [13,17]. The three breast cancer cell lines exhibited strikingly different requirements for exogenous adhesion support. Whereas MDA-MB-231 cells maintained adhesion without additional coating, SK-BR-3 cells required a serum-derived coating, and BT-474 cells failed to maintain adhesion under any of the conditions tested. These phenotypic differences were accompanied by cell-line-specific changes in the expression of adhesion-associated genes. Notably, FN1, which encodes fibronectin, was upregulated in adapted MDA-MB-231 cells, raising the possibility that endogenous ECM production contributes to their ability to maintain adhesion in the absence of exogenous coatings. However, the contribution of endogenous fibronectin to adhesion was not directly tested, and RNA-seq was performed on cells maintained on vitronectin-coated plates. Similarly, altered expression of adhesion-associated genes, including CEACAM6, NCAM2, and NRCAM, identifies potential mechanisms that may contribute to differences in substrate attachment. Functional studies will be required to determine whether these transcriptional changes directly influence the adhesion phenotypes observed during adaptation.

Adaptation was also accompanied by substantial transcriptional remodeling, including changes in pathways associated with cell-cycle regulation and metabolism. The nature of these changes differed among cell lines. In SK-BR-3 cells, adaptation was associated with reduced enrichment of E2F, G2M checkpoint, and MYC target gene sets, consistent with their reduced proliferation in 100% DM. In contrast, MDA-MB-231 and BT-474 cells exhibited distinct changes in growth-associated transcriptional programs, suggesting that reduced cell-cycle activity is not a universal response to adaptation. A more consistent feature was the alteration of metabolic pathways, particularly those involved in lipid and cholesterol metabolism. Cholesterol homeostasis gene sets were enriched in adapted SK-BR-3 and MDA-MB-231 cells, while expression of SREBF2 and its targets DHCR24 and FDFT1 increased across all three cell lines. These changes are consistent with a compensatory response to the reduced availability of serum-derived cholesterol and lipoproteins, potentially involving increased endogenous cholesterol synthesis and uptake [32]. However, direct measurements of cholesterol abundance and metabolic flux will be necessary to establish whether these transcriptional changes reflect altered cholesterol metabolism. Together, these findings suggest that adaptation to defined growth conditions is accompanied by substantial metabolic remodeling, with both shared and cell-line-specific responses.

These findings have implications for the use of conventional culture systems to investigate cancer metabolism and therapeutic vulnerabilities. Standard culture media supplemented with 10% fetal bovine serum contain an incompletely defined mixture of nutrients, lipids, and growth factors that may not reflect their availability in vivo. Previous studies have demonstrated that medium composition can influence metabolic dependencies, gene essentiality, and drug sensitivity [6,8]. Consequently, conventional serum-containing culture conditions may obscure metabolic vulnerabilities that emerge under more physiologically relevant nutrient conditions. The adaptation strategy described here provides an approach for establishing breast cancer cell populations capable of sustained growth in a defined nutrient environment. Such models could be useful for investigating metabolic dependencies through genetic screening or evaluating responses to metabolism-targeting therapies. Importantly, comparisons with cells maintained in conventional media will be necessary to determine how these altered culture conditions influence specific genetic and pharmacological vulnerabilities.

Several limitations should be considered when interpreting these findings. First, only three breast cancer cell lines were examined, with one representative of each molecular subtype, limiting the ability to generalize differences in adaptation capacity across subtypes. In addition, the cell lines were evaluated at different adaptation endpoints: SK-BR-3 and MDA-MB-231 cells were maintained in 100% DM, whereas BT-474 cells could only be maintained at a CM:DM ratio of 10:90 (1% FBS). The final culture conditions also differed in their substrate requirements. In particular, the serum-derived coating required by SK-BR-3 cells introduces undefined factors into an otherwise serum-free culture system, while MDA-MB-231 cells were analyzed on vitronectin-coated surfaces. Furthermore, because adaptation involved simultaneously reducing serum supplementation and changing basal medium composition, the contribution of individual medium components to the observed phenotypes cannot be distinguished. Prolonged adaptation may also reflect selection of preexisting cell populations, transcriptional reprogramming, or a combination of these processes. Finally, although HPLM more closely approximates the concentrations of circulating metabolites than conventional media, plasma composition does not necessarily reflect the nutrient availability or other environmental conditions experienced by cancer cells within tumors, either at the primary site or at metastatic sites. Additional studies incorporating tumor-relevant nutrient conditions and functional metabolic analyses will be necessary to establish how closely these adapted models reproduce cancer cell behavior in vivo.

In summary, we establish a stepwise approach for adapting breast cancer cells to growth in a more physiologically relevant, defined nutrient environment. Our findings highlight substantial differences in the capacity of individual cell lines to tolerate serum withdrawal and identify adhesion requirements and metabolic remodeling as important features of this process. Although further refinement is needed to better reproduce the tumor microenvironment, these adapted culture models provide a foundation for investigating how nutrient availability influences breast cancer cell biology and therapeutic vulnerabilities.

## Methods

### Cell lines and culture conditions

MDA-MB-231, SK-BR-3, and BT-474 cells were obtained from ATCC and maintained in CM prior to adaptation. Cells were cultured at 37°C in a humidified atmosphere containing 5% CO_2_. MDA-MB-231 cells were maintained in DMEM supplemented with 10% FBS and 1% penicillin-streptomycin, SK-BR-3 cells in McCoy’s 5A medium with the same supplements, and BT-474 cells in Hybri-Care medium supplemented with 10% FBS, 1% penicillin-streptomycin, and 1.5 g/L sodium bicarbonate. All cell lines were confirmed to be mycoplasma-negative prior to initiation of adaptation experiments.

### Defined media preparation

DM was prepared using HPLM supplemented with 1% penicillin-streptomycin, insulin-transferrin-selenium-ethanolamine (ITSE; 1:100 dilution of a 100X stock), hydrocortisone (0.5 µg/mL), human epidermal growth factor (EGF; 10 ng/mL), bovine serum albumin (BSA; 25 µM), and a 95:5 oleate:palmitate fatty acid mixture (100 µM total). To prepare the fatty acid component, BSA, oleic acid, and palmitic acid stock solutions were thawed at room temperature for 30 min. HPLM containing 1% penicillin-streptomycin was aliquoted into six 50-mL conical tubes (40 mL each) and supplemented with BSA, oleate, and palmitate at twice these final concentrations (53 µM, 202 µM, and 10.6 µM, respectively). These aliquots were incubated at 37°C with shaking for 1 hr to allow fatty acid complexing. In parallel, ITSE, hydrocortisone, and EGF were added to the remaining 260 mL of HPLM. Following incubation, the fatty acid-containing aliquots were recombined with the supplemented HPLM to generate the final DM formulation.

### Adaptation to defined medium

Cells were maintained in complete medium (CM) and passaged at 70 to 80% confluence. After one passage in CM, CM was progressively replaced with DM at CM:DM ratios of 75:25, 50:50, 40:60, 30:70, 20:80, and 10:90, followed by 100% DM, corresponding to 7.5, 5, 4, 3, 2, 1, and 0% FBS. Medium was mixed at the indicated ratio immediately before each medium change, and cells were maintained at each ratio for one to four passages before proceeding to the next. BT-474 cells received β-estradiol (0.5 nM) from the 40:60 ratio onward.

For each passage, cells were seeded at 3 x 10^6^ cells per 15-cm plate. At harvest, the culture medium and the PBS wash were collected and combined with the trypsinized cells, so that detached cells were retained in the count. For all three lines, medium was changed every 2 to 3 days, and at each change the removed medium was centrifuged, and any detached cells were returned to the plate with the fresh medium.

Once cells reached 100% DM, they were transferred to 6-cm plates carrying one of the five coating conditions described in Culture plate coating. Population doublings were calculated as log2(N/N0), where N is the total number of cells collected, and N0 is the number seeded, and cumulative population doublings were calculated by summing the values from each passage.

### Culture plate coating

To promote cell adhesion in 100% DM, 6-cm tissue culture plates were left uncoated or coated with FBS-containing medium, vitronectin (VN), fibronectin (FN), or a combination of VN and FN. For coating with FBS-containing medium, 2 mL of the corresponding cell line’s standard growth medium containing 10% FBS was added to each plate and incubated for 1 to 3 days at 37°C prior to use. Prior to seeding, medium was aspirated, plates were washed twice with PBS, and 100% DM was added.

For VN coated plates, VN was diluted in PBS to a final concentration of 0.5 µg/cm^2^ and added directly to 6-cm plates in a total volume of 2 mL. Plates were incubated for 2 hr at 37°C and subsequently stored at 4°C for 1 to 3 days before use. Prior to cell seeding, coated plates were washed twice with PBS and equilibrated with 100% DM. FN coated plates were prepared identically using FN at 1.0 µg/cm^2^. For dual coated plates, FN and VN were combined at final concentrations of 0.5 µg/cm^2^ each and applied using the same coating procedure.

### Immunofluorescence staining and imaging

Parental cells and BT-474 cells maintained in 10:90 CM:DM were seeded onto poly-L-lysine-coated coverslips. For 100% DM conditions, SK-BR-3 cells were seeded onto coverslips coated with FBS-containing medium, and MDA-MB-231 cells were seeded onto VN-coated coverslips. The following day, cells were fixed with 4% paraformaldehyde for 10 min at room temperature, washed with PBS, and permeabilized with 0.2% Triton X-100 for 5 min. Cells were blocked with 0.2% BSA and 2% normal goat serum (NGS) in PBS for 1 hr at room temperature, followed by incubation for 1 hr at room temperature with primary antibodies diluted in PBS containing 2% BSA. Primary antibodies included rabbit anti-HER2 (29D8, 0.945 µg/mL) for SK-BR-3 and BT-474 cells, and rabbit anti-vimentin (D21H3, 0.47 µg/mL) for MDA-MB-231 cells.

Cells were then washed three times in PBS and incubated for 1 hr in the dark with goat anti-rabbit Alexa Fluor 568 secondary antibody (4 µg/mL), Alexa Fluor 647-conjugated mouse anti-Ki-67 (B56, 1 µg/mL), Alexa Fluor 488-conjugated phalloidin (0.218 µg/mL), and DAPI (5 µg/mL). After three final 5 min PBS washes, coverslips were mounted onto glass slides with ProLong Gold Antifade Mountant.

Images were acquired on an Andor Dragonfly spinning-disk confocal microscope using 20x air (0.75 NA) and 63x oil immersion (1.40 NA) objectives. For quantitative analyses, 20x images were acquired as five-by-five montages with 50-plane z-stacks. Representative images were acquired using the 63x objective with 50-plane z-stacks. Fluorophores were excited with 405, 488, 561, and 637 nm lasers, and images were acquired using a Zyla sCMOS camera.

Z-stacks were processed by maximum-intensity projection and analyzed in CellProfiler (v4.2.8). Nuclei were segmented from the DAPI channel, and cell boundaries were delineated using the phalloidin channel. Mean Ki-67 intensity was measured for each nucleus, and mean HER2 or vimentin intensity was measured for each cell. Images flagged during segmentation for high nuclei counts or abnormally large cell areas were excluded.

Ki-67 positivity was determined in R (v4.4.1). Nuclei were classified as Ki-67-positive if their mean Ki-67 intensity exceeded the mean Ki-67 intensity measured in the corresponding cell line’s IgG control images, which were imaged and processed the same way. The percentage of Ki-67+ cells was calculated for each condition, with at least 333 cells quantified per condition. Bars show the mean percentage of Ki-67+ cells ± SEM. Box plots show the median, interquartile range (25th-75th percentiles), and whiskers extending to extreme values within 1.5 interquartile ranges of the quartiles. Pairwise comparisons were performed with two-sided Wilcoxon rank-sum tests on cell-level measurements, and p-values were adjusted within each plot using the Benjamini-Hochberg method.

### Bulk RNASeq

Total RNA was isolated with RNeasy spin columns. RNA sequencing of SK-BR-3 and MDA-MB-231 samples was performed by the Fred Hutch Genomics Shared Resource on an Illumina NovaSeq X Plus (paired-end 159-bp reads). BT-474 samples were sequenced by Azenta/Genewiz using poly(A)-selected libraries (150-bp paired-end reads). Reads were processed using nf-core/rnaseq (v26.0) [33], including trimming, alignment to GRCh38 using GENCODE v50 annotation, and gene-level quantification with Salmon and tximport.

Downstream analyses were performed in R. Differential gene expression was analyzed using DESeq2 (v1.46.0), comparing three biological replicates of parental cells with three replicates of adapted cells for each cell line (100% DM for SK-BR-3 and MDA-MB-231, 10:90 CM:DM for BT-474). Genes were called differentially expressed if they had a Benjamini-Hochberg adjusted p-value < 0.05 and an absolute log2 fold change > 1.

Volcano plots and GSEA used protein-coding genes with at least 10 total counts across the six samples for each cell line. For GSEA, genes were ranked by the DESeq2 Wald statistic and analyzed against the MSigDB Hallmark collection (preranked; clusterProfiler v4.14.6, fgsea v1.32.4, msigdbr v26.1.0; gene sets of 10 to 500 genes). The 10 Hallmark gene sets with the lowest adjusted p-value are shown.

To examine adhesion-associated gene expression, all MSigDB C5 GO Biological Process gene sets with “adhesion” in their name were retrieved with msigdbr, and the genes across these 70 gene sets were combined (∼1,400 unique genes) to define the adhesion gene list. Adhesion genes were extracted from the DESeq2 results using the same filters as above. The complete DESeq2 results for each comparison, together with the adhesion gene list, are provided in Supplementary Table S1. For Figure 4J, the 12 adhesion genes per line with the largest absolute Wald statistics were displayed as log2 fold change bars, with Benjamini-Hochberg-adjusted p-values indicated with asterisks.

## Acknowledgements

We thank members of the Alvarez lab for helpful discussions. We thank Lucas Sullivan and Oliver Newsom for advice on defined medium formulations. This research was supported by National Cancer Institute grant R01CA292658 (J.V.A), through funding from Seattle Translational Tumor Research (STTR), by the Cellular Imaging and Genomics and Bioinformatics Shared Resources of the Fred Hutch/University of Washington Cancer Consortium (P30 CA015704), and by startup funds from Fred Hutch Cancer Center (J.V.A).

## Supplementary Figure 1

**Supplemental Figure 1.**
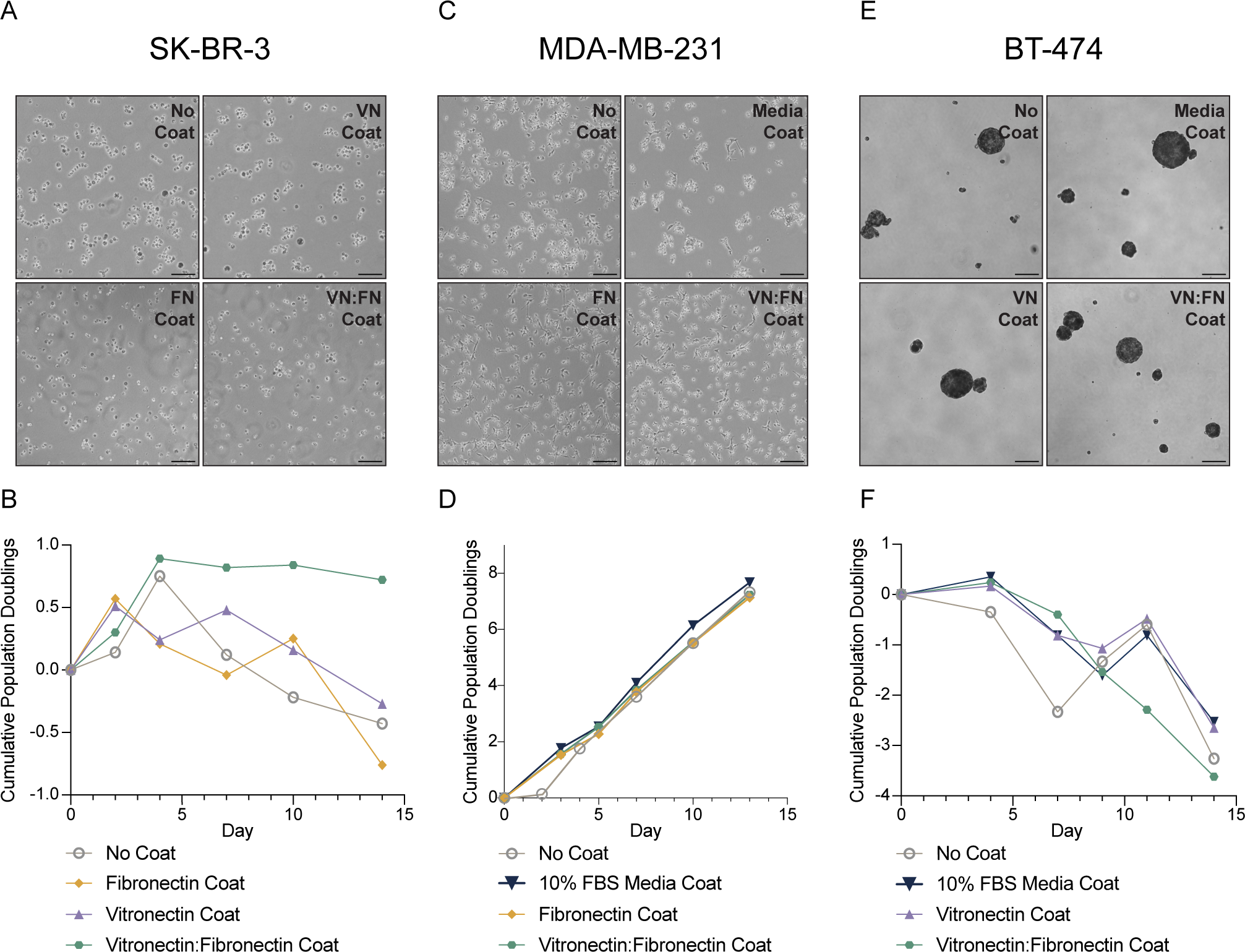
Coating conditions tested for maintenance of breast cancer cells in 100% DM. **(A, C, E)** Representative phase-contrast images of SK-BR-3 **(A)**, MDA-MB-231 **(C)**, and BT-474 **(E)** cells maintained in 100% DM on uncoated plates or plates coated with vitronectin (VN; 0.5 µg/cm²), fibronectin (FN; 1.0 µg/cm²), combined FN/VN (0.5 µg/cm² each), or FBS-derived coating, as indicated. **(B, D, F)** Cumulative population doublings of SK-BR-3 **(B)**, MDA-MB-231 **(D)**, and BT-474 **(F)** cells cultured in 100% DM on the coating conditions shown above. The coating conditions selected for subsequent maintenance of SK-BR-3 and MDA-MB-231 cells are shown in Figures 1 and 2, respectively. FN-coated plates were used to evaluate BT-474 cells following transfer to 100% DM, although no coating condition supported sustained proliferation. Scale bars, 150 µm.

## Supplemental Tables

**Supplementary Table S1. Differential gene expression results for adapted breast cancer cells.** Complete DESeq2 results for each parental-versus-adapted comparison: SK-BR-3 (parental vs 100% DM on the 10% FBS media coating), MDA-MB-231 (parental vs 100% DM on VN coating), and BT-474 (parental vs 10:90 CM:DM, 1% FBS). Each sheet lists every gene with at least one read, with baseMean, log2 fold change, standard error, Wald statistic, P value, Benjamini-Hochberg-adjusted P value, whether the gene meets the differential-expression thresholds (adjusted P < 0.05, |log2 fold change| > 1), and the direction of change. Column definitions, processing details, and software versions are provided in the README sheet.

**Supplementary Table S2. Reagents and resources used in this study.** Cell lines, antibodies, defined-medium components, base-media components, coating reagents, and immunofluorescence reagents are listed with their sources and catalog identifiers.

