## Supplemental Table S2 for "Adaptation of breast cancer cells to serum-free defined medium reveals changes in proliferation, adhesion, and lipid metabolism"

**Supplementary Table S2. Reagents and resources used in this study.**

| **REAGENT or RESOURCE** | **SOURCE** | **IDENTIFIER** |
| --- | --- | --- |
| **Cell Lines** | | |
| MDA-MB-231 | ATCC | Cat. #: HTB-27 |
| SKBR-3 | ATCC | Cat. #: HTB-30 |
| BT-474 | ATCC | Cat. #: HTB-20 |
| **Antibodies** | | |
| HER2/ErbB2 (29D8) Rabbit mAb | Cell Signaling Technology | Cat. #: 2165S  RRID: AB_10692490 |
| Vimentin (D21H3) Rabbit mAb | Cell Signaling Technology | Cat. #: 5741  RRID: AB_10695459 |
| Ki-67, Alexa Fluor 647 Mouse mAb, clone B56 | BD Pharmingen (BD Biosciences) | Cat. #: 558615  RRID: AB_647130 |
| Rabbit (DA1E) mAb IgG XP® Isotype Control | Cell Signaling Technology | Cat. #: 3900S  RRID: AB_1550038 |
| Goat anti-Rabbit IgG (H+L), Alexa Fluor 568 (secondary) | Thermo Fisher Scientific (Invitrogen) | Cat. #: A-11036  RRID: AB_10563566 |
| **Chemicals, Peptides, and Recombinant Proteins** | | |
| *Defined Media (DM) Components* | | |
| Human Plasma-Like Medium (HPLM) | Gibco | Cat. #: A4899101 |
| Penicillin-Streptomycin | Gibco | Cat. #: 15140122 |
| Oleic Acid | Sigma-Aldrich | Cat. #: O1008-1G |
| Palmitic Acid | Sigma-Aldrich | Cat. #: P0500-10G |
| Bovine Serum Albumin (BSA), fatty-acid carrier stock | Sigma-Aldrich | Cat. #: 3117057001 |
| Insulin-Transferrin-Selenium-Ethanolamine (ITSE) | Biogems International | Cat. #: 50-152-8628 |
| Hydrocortisone | Sigma-Aldrich | Cat. #: H0396-100MG |
| Human EGF | PeproTech | Cat. #: AF-100-15-100UG |
| *Base (10% FBS) Media Components* | | |
| DMEM, Dulbecco's (500 mL) | Gibco | Cat. #: 11965092 |
| McCoy's 5A (Modified) Medium | Gibco (Thermo Fisher Scientific) | Cat. #: 16600082 |
| Hybri-Care Medium | ATCC | Cat. #: 46-X |
| Fetal Bovine Serum (FBS), Value FBS (500 mL) | Gibco | Cat. #: A5256701 |
| Sodium Bicarbonate | Sigma-Aldrich | Cat. #: S5761-500G |
| β-Estradiol (BT474 media only; added per-row, not to bottle) | Sigma-Aldrich (Millipore Sigma) | Cat. #: E2758-250MG |
| *Coating Reagents* | | |
| Human Vitronectin | Sigma-Aldrich (Millipore Sigma) | Cat. #: SRP3186-250UG |
| Bovine Fibronectin | Sigma-Aldrich (Millipore Sigma) | Cat. #: F4759-2MG |
| *Immunofluorescence (IF) Reagents* | | |
| Poly-L-Lysine Solution, 100 mL | Cell Biologics | Cat. #: 6951 |
| Triton X-100 (permeabilization) | Sigma-Aldrich | Cat. #: T8787-250mL |
| Normal Goat Serum (NGS, blocking buffer) | Gibco (Thermo Fisher Scientific) | Cat. #: PCN5000 |
| Bovine Serum Albumin (BSA, IF blocking buffer) | Sigma-Aldrich | Cat. #: B4287-25G |
| Phalloidin, Alexa Fluor 488 conjugate | Invitrogen (Thermo Fisher Scientific) | Cat. #: A12379 |
| DAPI (4',6-Diamidino-2-Phenylindole, Dihydrochloride) | Invitrogen (Thermo Fisher Scientific) | Cat. #: D1306 |
| ProLong Gold Antifade Mountant | Invitrogen (Thermo Fisher Scientific) | Cat. #: P36930 |
